# Prevalence of Soil Transmitted Helminthes (STHs)Among Pupils of Community Primary Schools in Nkpor and Mgbodohia, Obio/Akpor Local Government Area, Rivers State, Nigeria

**DOI:** 10.1101/464008

**Authors:** LeBari Barine Gboeloh, Ike-Ihunwo Chikaire Ndamzi

**Affiliations:** Department of Biology, Ignatius Ajuru University of Education, Port Harcourt, Nigeria; Rivers State College of Health Science and Technology, Port Harcourt

**Keywords:** Prevalence, Intensity, Soil-transmitted helminth, Nkpor, Mgbodohia

## Abstract

Soil transmitted helminthes (STHs) are common public health concern among children in Sub saharan Africa. A study to determine the prevalence and intensity of these parasites among pupils of two primary schools in Nkpor and Mgbodohia communities, Obio/Akpor Local Government Area, Rivers State, Nigeria was conducted. The formo-ether concentration technique was used to concentrate and separate the eggs and cysts from the faeces. Out of 107 pupils (56males and 51 females) investigated, 81 (75.7 %) were positive for at least one gastrointestinal helminth. The parasites identified included *Ascaris lumbricoide*(58.0%), Hookworms(28.4%), *Trichirus trichiura*(13.6%) and *A.lumbricoide* + *T. trichiura* (6.3%). *Ascaris lumbricoide* was significantly (P<0.05) higher in prevalence than other parasites. Although more females (54.3%) were infected than males (45.7%), there was no significance(P>0.05) difference in the prevalence in relation to sex. Of the 44 males infected, 24(54.5%), 5 (11.4%), 13(29.5%) and 2(4.5%) haboured *A. lumbricoide, T. trichiura*, Hookworms and mixed infection (*A. lumbricoide + T. trichiura*) repectively. Out of the 37 females infected, 21 (56.8%), 3(8.1%), 10(27.0%) and 3(8.1%) haboured *A. lumbricoide, T. trichiura*, Hookworms and mixed infection (*A. lumbricoide + T. trichiura*) repectively. There was no significance difference in the prevalence of *A. lumbricoide* and Hookworms between males (54.5%) and females (56.8%). There was significant difference in prevalence among two major age groups (5-10years-45% and 11-15years-41.9%). Children within the age group of 16-20years had the least infection (9.9%). The intensities of *A.lumbricoide, T.trichiura* and hookworm were 246.5, 107.5 and 187 Epg respectively. The intensity of *A. lumbricoide* was significantly difference than other parasites identified.Soil transmited helminthes remain a public health concern among children in the study area. Provision of portable water, toilet facilities, good education on the epidemiology of STHs and regular de-worming will enhance control measures.

## 1.0 Introduction

*Soil*-transmitted helminths are a group of gastrointestinal parasites that are transmitted to human through contaminated soil. These parasites have their developmental stages in the soil and transform into infective stage (Chukwuma *et al*, 2009). The eggs are passed out in faeces of infected persons into the environment, hence contaminating the soil in areas with poor sanitation especially where there are no modern toilet facilities. The infections are caused by different species of parasitic worms. These parasites are majorly nematodes and include round worm (*Ascaris lumbricoides*), whipworm (*Trichuris trichiura*), and the species of hookworm (*Anclostoma duodenale* and *Necator americanus*) (Hotez *et al*, 2007). Helmithiasis (Ascariasis, trichuriasis and hookworm disease) caused by these parasites are the most prevalent in developing countries of the world possibly because of poor sanitary condition coupled with poverty and other environmental factors that encourage the thriving of the parasites.

These infections constitute a global public health concern especially in sub Saharan Africa where poverty and poor sanitary condition are endemic. The World Health Organization recorded that an estimated 1.5 billion people or 24% of global population are infected with soil-transmitted helminths worldwide(WHO, 2007a), out of which an estimated 300 million cases result in severe morbidity. However, the morbidity is associated with heavy worm burdens(Hotez *et al*, 2003). In 2010, about 438.9 million people were infected with hookworm, 819.0 million with *A. lumbricoides* and 464.6 million with *T. trichiura*(NCDC, 2007; Rachel *et al*, 2007; Rachel *et al*, 2002). Ascariasis caused an estimated 2,824 deaths large percentage of which occurred in Asia (Rachel *et al*, 2007). In 2016, one-third of an estimated three billion people living on less than two US dollars per day in developing countries of the world including Sub-Saharan Africa were at risk of at least one soil transmitted helminthic infection (Ngonjo *et al*,2007; Hotez *et al*, 2007).

In sub-Saharan Africa, about 866 million people were infected by STH in 2012. Hookworm, *A. lumbricoides*, and *T. trichiura* accounted for 117 million (13.6%), 117 million (13.6%), and 100.8 million (11.6%) of the total infection respectively(WHO, 2007b). Although the infections are reportedly prevalent in rural areas, people living in slums and scattered settlements in urban and peri-urban areas are also at risk (Crompton and Savioli, 1993; Salawu and Ughele, 2015).

School children are the most vulnerable group at risk of STH due to the habit of walking and playing barefoot, poor nutrition and poor awareness or education on the transmission pattern of the parasites (Crompton, 1989), lack of clean water, inadequate healthcare and poor personal hygiene (NCDC, 2007; Crompton,1989; Asaolu *et al*, 2002). The infections impact heavily on the physical growth and cognitive development of infected children(Stollzfus *et al*, 1996; Miguel and Kremer, 2004).

Globally, an estimated 267 million pre-school-age children and over 568 million school-age children live in endemic areas and are in need of treatment and preventive interventions WHO (2007b). These infections cause a wide range of abdominal complications, iron-deficiency anaemia and dysentery syndrome in children WHO (1998). WHO (2018) reported that STH weaken the nutritional status of the infected individuals in many ways including feeding on host tissues and blood leading to loss of iron and protein (anaemia) as in the case of hookworm, increased malabsorption of nutrients and competition for vitamin *A. Trichiura* trichiura causes dysentery and diarrhea while others have been implicated in loss of appetite which invariable reduce intake of nutrients causing poor physical growth.

To cub this trend, in 2001, delegates at the World Health Assembly unanimously endorsed resolution WHA54.19 which urged member countries in endemic regions to give serious attention to the tackling of STHs with the view to control morbidity through periodic dewormimg of school children and other persons at risk of the disease. Again, the resolution also recommended health education and provision of adequate sanitation and personal hygiene as intervention measures for the control of STHs, and recommended albendazale and mebendazole as drugs of choice for the treatment of infected persons WHO (2018).

In Nigeria, several studies have been conducted on the prevalence of STH in different regions of the country (WHO 2018; Ofoezie *et el,* 1996; Salawu and Ughele, 2015). De-worming exercise is intermittently embarked upon by individuals, Non-Governmental organizations and at times Government without background data(Odinaka *et al,* 2005; Ekpo *et al*, 2008) or baseline data upon which such program should be initiated violating the WHO’s recommendation for a baseline analysis to be conducted on the prevalence of STH before initiation of control program (Odu *et al*, 2011, WHO, 2012). In the study area, there is no available documented study with regard to the prevalence of these parasites among school children, hence it becomes pertinent to conduct this study with the view to establishing a baseline data for policy makers and research minded scholars.

## 2.0 MATERIALS AND METHODS

### 2.1 Study Area

Nkpor and Mgbodohia are two communities in Obio/Akpor Local Government Area. The communities have common boundaries and share common facilities. They are clans within Rumuolumeni kingdom. The communities are host to several multinational oil companies including Agip oil company and Eni group that have several subsidiaries. The Local Government Area is basically a low land area, about 30 meters above sea level. It covers about 100sq mi (260 km ^2^); and due to high rain fall, the soil consists of sandy or sandy loam(Eludoyin *et al*, 2011).The vegetation consist of light rain forest and thick mangrove forest. According to the 2006 Census, it has a population of 464,789(NPC 2006). It is located between latitudes 4°45′N and 4°60′N and longitudes 6°50′E and 8°00′E. Covering around 100 sq mi, Obio-Akpor is a lowland area with an average elevation below 30 meters above sea level.

### 2.2 Study Population

The study subjects were primary school pupils between the age of 5 to 16 years. The only two public primary schools in the area (Community Primary school, Mgbodohia and Community Primary School, Nkpor) were selected for the study. The study was conducted between Novemeber 2017 and January, 2018.

### 2.3 Sample Collection

A total of 150 pupils were randomly selected for study and the method of Gboeloh and Elele (2013) was adopted in the collection of faecal samples. Well labeled plastic specimen bottles containing 10% formaldehyde and wooden scapula were distributed to the pupils, for collection of their faecal sample. The pupils were properly instructed on how to carefully collect their faecal samples into the specimen bottles. The samples were transported to the Research Laboratory, Department of Biology, Ignatius Ajuru University of Education for parasitological examination. Furthermore, demographic information on the age and sex of participants were collected from the school record, the consistency of the stool (formed, soft, semi-soft and watery) were recorded for each subject on a recording format.

### 2.4 Parasitological Examination

Parasitological examination of the faecal samples was done using formo-ether concentration technique (Cheesbrough, 2005) to concentrate the parasites. About 1g of each faecal sample was thoroughly mixed in a test tube containing 4ml of 10% formol water and sieved with cotton gauze into centrifugal tube containing another 4ml of formol water. 3ml of diethyl ether was then added and thoroughly mixed. The mixture was then centrifuged for one minute at 3000rpm. The sediments were transferred onto a microscopic slide after the supernatant was discarded. Parasites eggs and cyst were observed and identified using X10 and X40 magnification of the microscope.

### 2.5 Ethical Consideration

The head teachers and Parent Teachers Association (PTA) of the selected schools were formally contacted in writing with regard to the research work. This was followed by official visit during which detail explaination on the essence of the research work made to the management of the schools and PTA. Further, the researchers interacted with the pupils at the early morning assembly ground during which orientation was given to the pupils about the research work. Subsequent upon this, written informed consent was also obtained from the Head Teachers and Parent Teacher Associations (PTA) of the schools investigated. Another approval was also given to the work by the Ethics Committee, Ignatius Ajuru University of Education, Port Harcourt.

### 2.6 Data Analysis

The prevalence rate (Pr) was determined by dividing the number of infected pupils (Ni)by the total number of pupils examined (Ne) multiplied by 100.

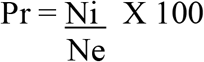

Parasite intensity (Pi) among the pupils was calculated as the number of eggs (Ng) per gram of faeces (Fg) while the parasite load was expressed as the mean number of eggs of each parasite species per gram of faeces of each pupil.

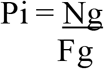

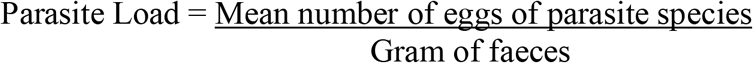

Statistical analysis of the data was done using SPSS (version 16.0). Chi-square test was also used to compare difference among variables at significance level of p <0.05.

## 3.0 Results

Out of the 150 pupils that received specimen bottles, only 107 returned their faecal samples. Of the 107 samples (56 males and 51 females) received and examined, 81 (75.7 %) were positive for at least one helminthic infection (Fig. 1.0). Of the 81 positive samples, 44 (54.3%) were females while 37(45.7%) were males. Of the 44 males infected, 24(54.5%), 5 (11.4%), 13(29.5%) and 2(4.5%) haboured *A. lumbricoide, T. trichiura*, Hookworms and mixed infection (*A. lumbricoide + T. trichiura*) repectively (Table 1.0). Similarly, of the 37 females positive for helminth parasites, 21 (56.8%), 3(8.1%), 10(27.0%) and 3(8.1%) haboured *A. lumbricoide, T. trichiura*, Hookworms and mixed infection (*A. lumbricoide + T. trichiura*) repectively(Table 1.0). There was no significance in the prevalence of *A. lumbricoide* and Hookworms between males (54.5%) and females (56.8%). However, there was statistical significance in the prevalence of *T. trichiura* between males (11.4%) and females (8.1%). Similarly, there was a statistical significance in prevalence of mixed infection between males (4.5%) and females (8.1%).

**Fig. 1.0:**
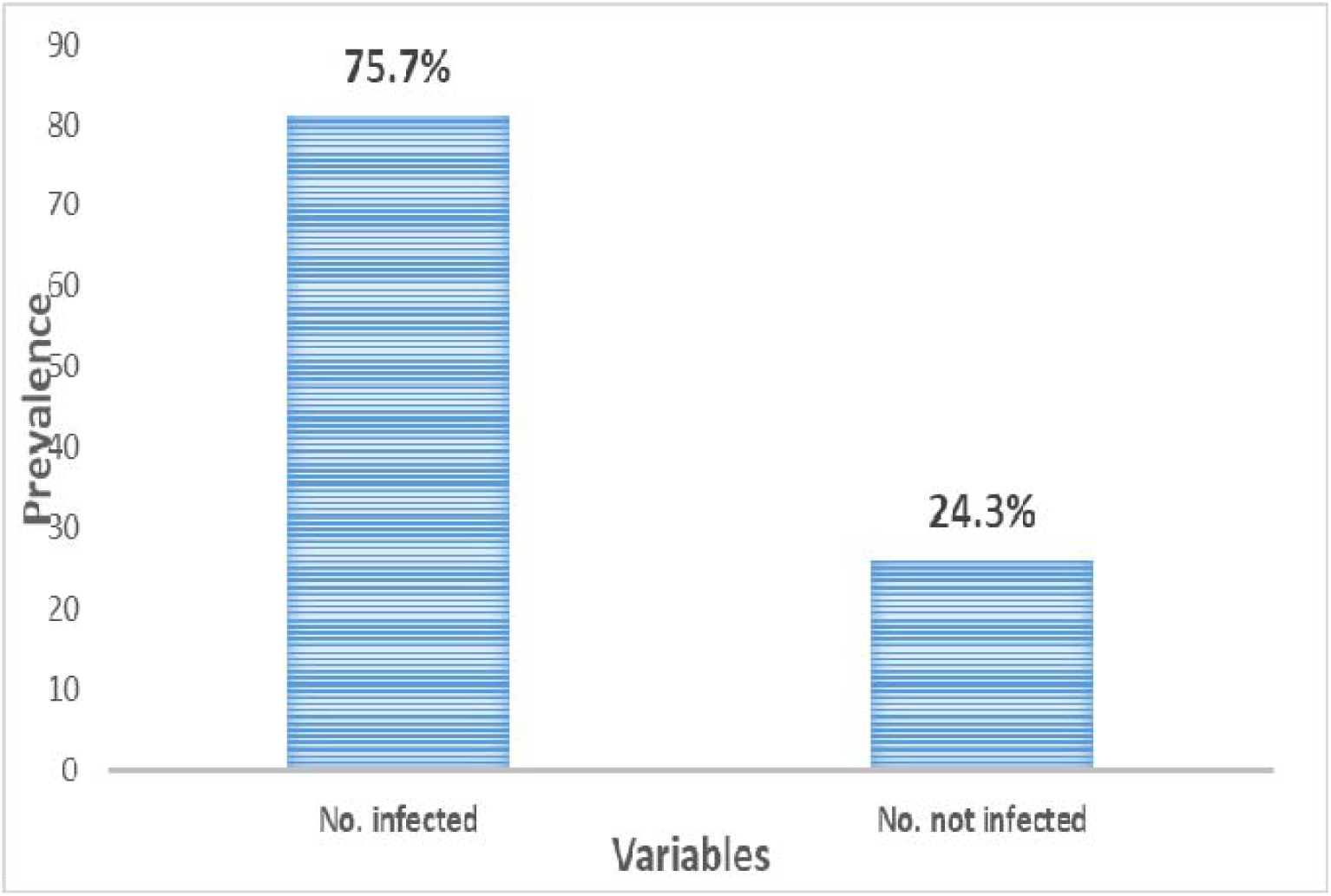
Overall prevalence of Soil Transmitted Helminthes (STH) among primary school pupils in Akpor and Mgbodohia

**Fig. 2.0:**
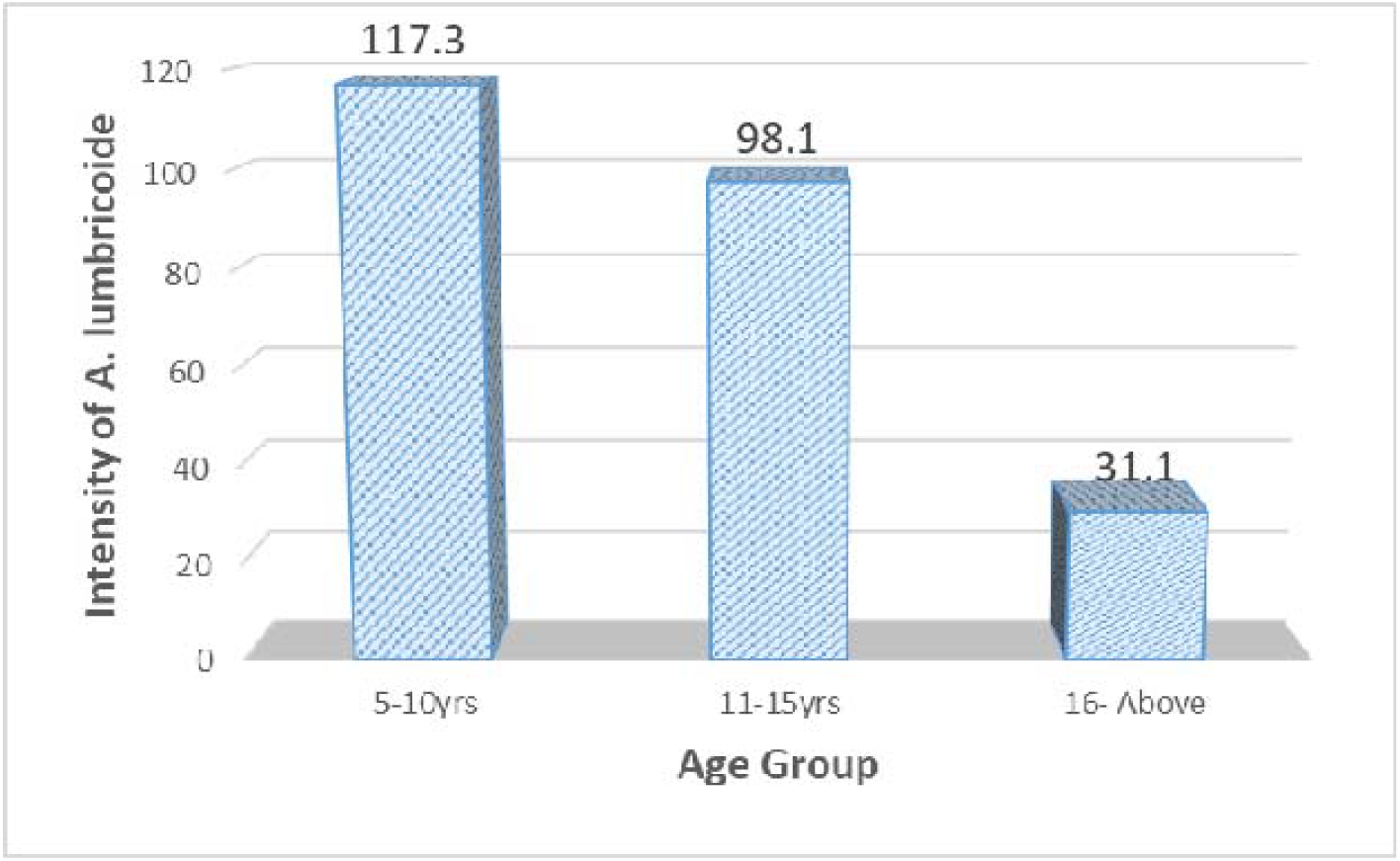
Intensity of *A. lumbricoide* in relation to age

**Fig. 3.0:**
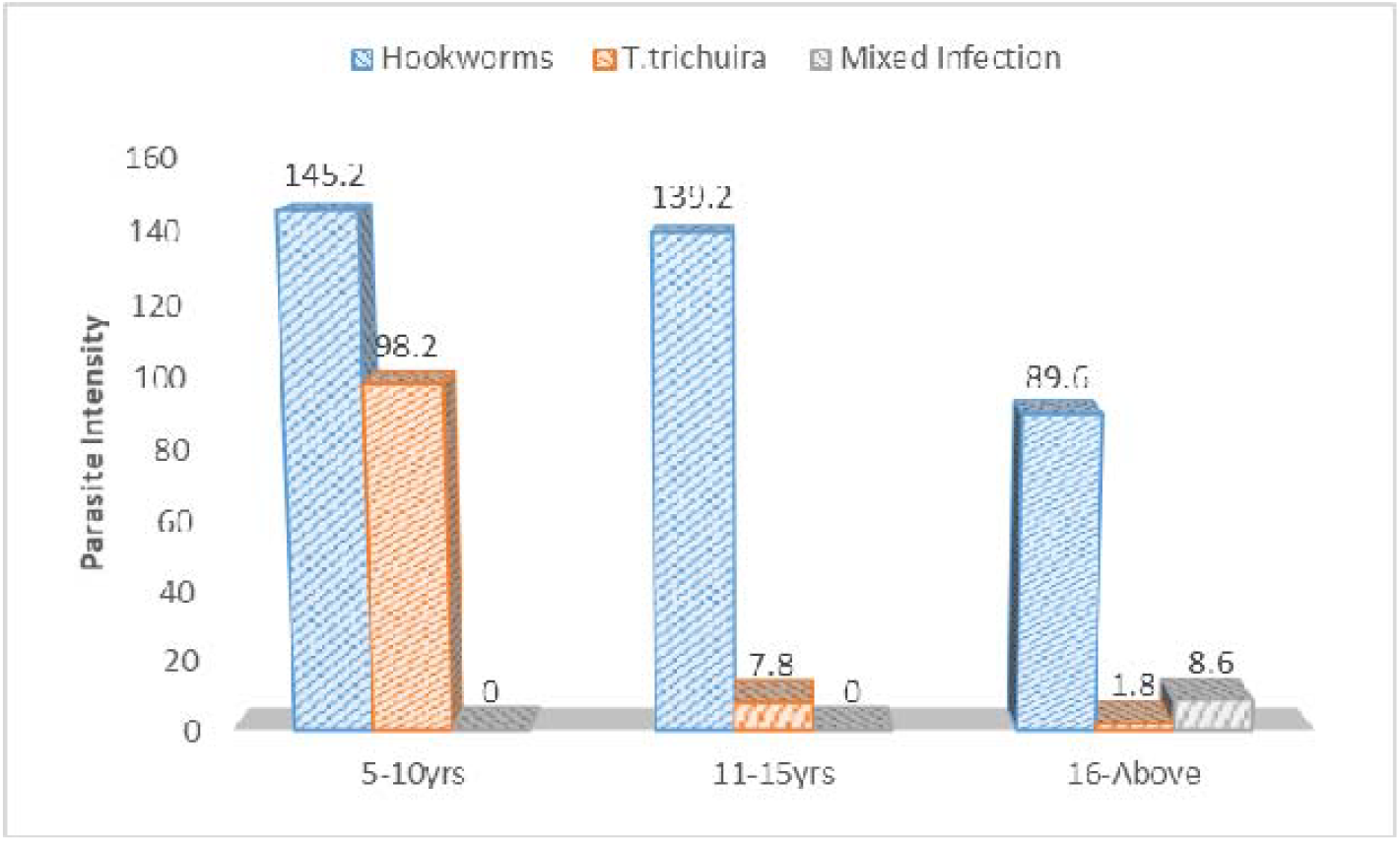
Intensity of Hookworms, *T. trichiura* and mixed infection

**Table 1.0:** Prevalence of helminth parasite in relation to sex (n= 107)

| Sex | No.<br>examined | Species of helminthes |  |  |  |  | Total infected (%) |
| --- | --- | --- | --- | --- | --- | --- | --- |
|  |  | *A.lumbricoide*<br>No. infected (%) | *T. trichiura*<br>No. infected(%) | Hookworms<br>No. infected(%) | Mixed Infection<br>(*A. Lumbricoide* +<br>*T. trichiura*) |  |  |
| Male | 56 | 24 (54.5) | 5(11.4) | 13(29.5) | 2 (4.5) |  | 44(78.6) |
| Female | 51 | 21(56.8) | 3(8.1) | 10 (27.0) | 3(8.1) |  | 37(72.5) |

The results indicated that there was no significance difference (p> 0.05) in the prevalence of the parasites between the two major age groups, 5-10years (45.7% and 11-15years (43.0%). However, children that were above 16years old had the least infection (12.3%). Similarly, *A. lumbricoide* had no significance prevalence across age group (Table 2.0). No significance difference in prevalence of Hookworms between age group 5-10years (27.0%) and age group 11-15years (32.4%). However, a statistical difference was observed in the prevalence of mixed infection (*A. lumbricoide* + *T. trichiura*) among children above 16years old and the rest of the age groups (Table 2.0). Out of the 81 children positive for STH, 47 had *Ascaris lumbricoide*, 23had Hookworm, 11 had *Trichirus trichiura* while 5 had mixed infection (*A.lumbricoide* + *T. trichiura*) representing 58.0%, 28.4%, 13.6% and 6.2% of the total worm burden respectively (Table 2). The infection was statistically significance for *A. lumbricoide* with Nkpor (53.2%) higher than Mgbuodohia (46.8%). There was a significance difference (p< 0.05) in prevalence between Nkpor (59.3%) and Mgboudohia (40.7%). The prevalence of hookworms (28.4%) in the study area was not significance. However, there was significance difference in the prevalence of hookworm between the two schools; Nkpor (60.9%) and Mgboudohia (39. 1%). In all, statistical significance difference (p< 0.05) was observed in the prevalence of *T. trichiura*, hookworms and mixed infection (*A. lumbricoide* and *T. trichiura*) between schools in the study area(Table 2).

**Table 2.0:**
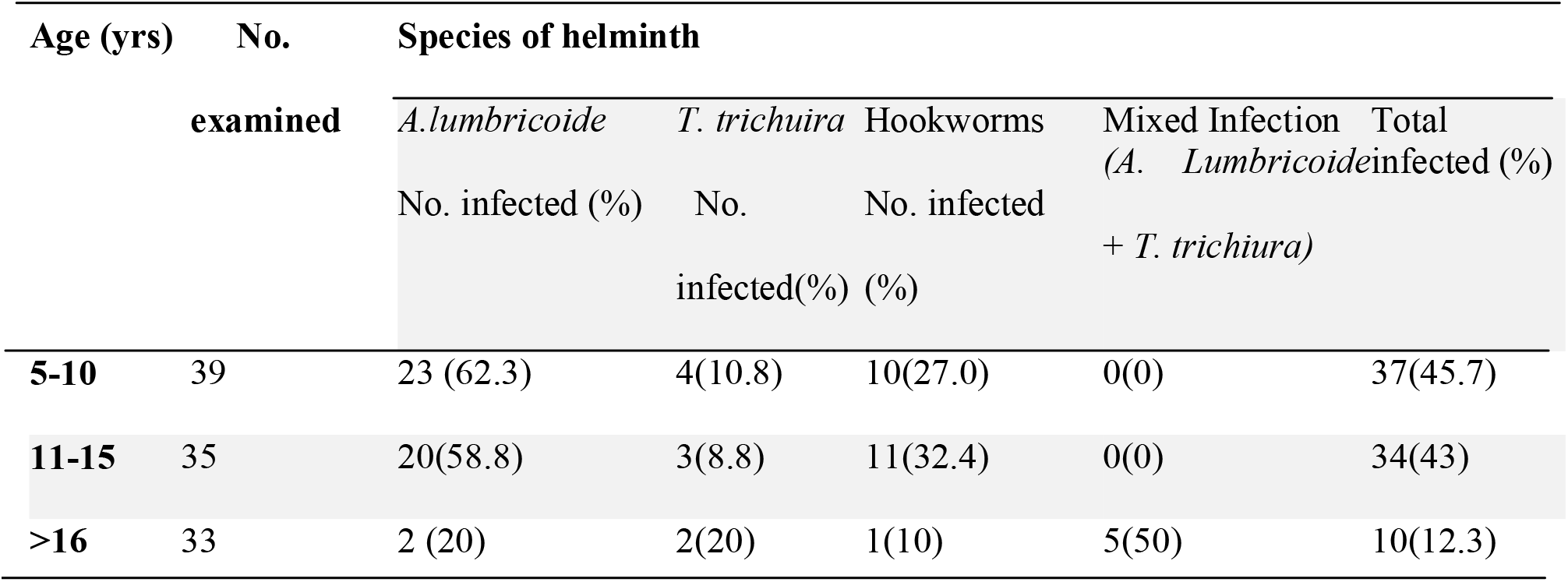
Prevalence of helminth parasites in relation to age (n=107)

| Age (yrs) | No.<br>examined | Species of helminth |  |  |  |  | Total infected (%) |
| --- | --- | --- | --- | --- | --- | --- | --- |
|  |  | *A.lumbricoide*<br>No. infected (%) | *T. trichiura*<br>No.<br>infected(%) | Hookworms<br>No. infected (%) | Mixed Infection<br>(*A. Lumbricoide* +<br>*T. trichiura*) |  |  |
| 5-10 | 39 | 23 (62.3) | 4(10.8) | 10(27.0) | 0(0) |  | 37(45.7) |
| 11-15 | 35 | 20(58.8) | 3(8.8) | 11(32.4) | 0(0) |  | 34(43) |
| >16 | 33 | 2 (20) | 2(20) | 1(10) | 5(50) |  | 10(12.3) |

### Parasite Intensity

The intensity of parasites was expressed as the number of eggs of specific parasite per gram of faeces. Among pupils of community primary school Mgbodohia, the intensity was 216.8, 188.7, 172.2 and 8.6 for *A. lumbricoide, T. trichiura*, Hookworm and Mixed infection (*A. lumbricoide and T. trichiura*) respectively. For community primary school, Nkpor, the intensity included 276.8, 26.8, 201.8 for *A. lumbricoide, T. trichiura* and Hookworm respectively. There was no statistically significance difference (P>0.05) in the intensity of *A. lumbricoide* and Hookworm in pupils of both schools. However, a significance difference (p <0.05) existed in the intensity of *T. trichiura* and mixed infection (*A. lumbricoide, T. trichiura)* among pupils of both schools (Table 3). The intensity of *A. lubricoide* was higher in children in the age group of 5-10years and lower in nchildren above 16years. Similar trend was observed in the intensity of hookworm and *T. trichiura.*

**Table 3.0:**
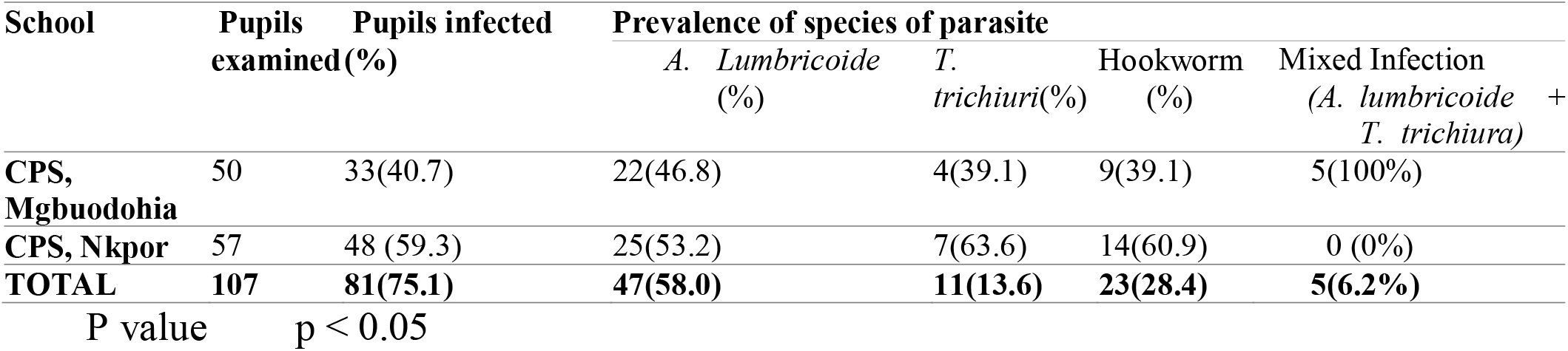
Prevalence of Soil Transmitted Helminthes (STH) in relation to species of parasites (n= 107)

**Table 3.0:** Intensity(epg) of Soil Transmitted Helminthes among primary school pupils in Nkpor and Mgboudohia

| School | Species of helminthes |  |  |  |
| --- | --- | --- | --- | --- |
|  | *A. lumbricoide* | *T. trichiura* | Hookworms | Mixed Infection (*A. Lumbricoide* + *T. trichiura*) |
| **CPS, Mgbuodohia** | 216.8 | 188.7 | 172.2 | 8.6 |
| **CPS, Nkpor** | 276.2 | 26.8 | 201.8 | 0 |
| **TOTAL** | 246.5 | 107.8 | 187 | 4.3 |

## 4.0 Discussion

The occurrence of soil transmitted helminthes among primary school children in Mgbodohia and Nkpor communities was investigated. A statistically significant proportion (75.7%) of the population in the study area haboured at least a helminth parasite, an indication that the prevalence rate of these parasites is high in the study area and still remains a public health concern among children (Nwosu, 1981; Asaolu *et al*, 2002). Similar trend has been reported among school age children in various parts of Nigeria (Ukpai and Ugwu, 2003; Egwunyenga and 2005; Ugbomoiko *et al*, 2006; Adeoye *et al*, 2007; Obiukwu *et al*, 2008; Sowemimo *et al,* 2011; Okolo and John, 2008; Aisien *et al,* 2002). However, the high prevalence rate recorded in this study is higher than the 59.2% recorded by (Salawu and Ughele, 2015) among school-age children in Ife, Osun state and the 30.3% reported by Ugbomoiko and Ofoezie (2007) among school children in Amaruru community, Imo State, Southeast Nigeria. It is also higher than the 24.6% observed by Sowemimo and Asaolu (2011) among among rural Fulani children in Vom, Plateau State. The result is also higher than the 58.0% reported by Bangural *et al* (2016) among primary school children in Bo district of Sierra Leone, the 51.5% repoted by Bopda *et al* (2016) among adults of Akonolinga health district of Cameroon and the 19.3% recorded by Adu-Gyasi *et al* (2018) in the middle-belt of Ghana. The result was however lower than the 90% prevalence rate recorded among high risk area in India by Salma and Azam (2007) and the 75.6% recorded among primary school children in another part of India by Ganguly *et al* (2017). The high prevalence recorded in this study could be attributed to lack of toilet facilities and portable drinking water (Ugbomoiko and Ofoezie. 2017) in the area. The poor socio-economic status and occupation of the parents may also be responsible for the high prevalence of the parasites among these children (Salawu and Ughele, 2015; Ugbomoiko and Ofoezie, 2017). Furthermore, the high prevalence rate could also be attributed to the fact that the studied primary schools are located in rural communities. The two communities are located at the periphery of Port Harcourt metropolis and had no urban flavor. The communities lack social infractures such as safe drinking water, good road and electricity. Ojurongbe *et al* (2014) reported that rural communities are characterized by high poverty rate, poor infrastructural development, poor sanitation and poor hygiene, factors that enhance the transmission of helminth parasites.

Most of the pupils (58%) haboured *Ascaris lumbricoide*. The prevalence rate of *A. lumbricoide* observed in this study is higher than the 21.8% and 13.1% reported by Asaolu *et al* (2002) in Ifewara and Ajebandele respectively, 4% in Edo (Oguanya *et al*, 2012), 18.8% in Adamawa (Shitta and Akogun, 2017), 1.5% in Ghana (Adu-Gyasi *et al*, 2018) and 18.8% in Cameroon (Bopda *et al*, 2016). The high prevalence reported in this study is also comparatively higher than the 44.8%, 13.1% qnd 17.7% reported by Salawu and Ughele (2015), Asaolu *et al* (2002) and Bangural *et al* (2016) respectively. The parasite also occured in all age groups investigated confirming the findings of Bangural *et al* (2016) and Egwunyenga and Ataikiru, (2005). These authors recorded the prevalence of *A. lumbricoide* in all age groups investigated.

The high prevalence rate among children of lower age group recorded in this study is contrary to the findings of Salawu and Ughele (2015) and Sowemimo and Asaolu (2011). The researchers recorded a the lowest prevalence among lower age groups below 10years. The difference in results could be attributed to the differences in study area and the attitude of parents towards their children. Our study was conducted in rural communities were little children are allowed to crawl on the ground and play indiscremately on the ground and sometime bare footed. Moreso, they eat with unwashed or inproperly washed hands. The lower prevalence among children above 16years old is in consonance with the report of other authors (Salawu and Ughele, 2015; Sowemimo and Asaolu, 2011; Bangural *et al*, 2016). The lower prevalence among this age group could be as a result of the fact that as age increases, the children become more conscious of personal hygiene and avoid playing with soil and unhygienic places (Salawu and Ughele, 2015; Adanyi *et al*, 2011). They have less contact with soil as they are old enough to control their playing habit and are more conscious of their personal hygiene (Odinaka *et al*, 2005). Similar observation has been recorded by Ukpai and Ugwu (2003) and Obiukwu *et al* (2008).

The prevalence of Hookworms (28.4%) in the present study is lower than the 94.2% recorded among Primary School Children in a Amaruru Community in Imo State by Odinaka *et al* (2005). It is also lower than the 33.3% recorded by Gboeloh and Elele (2013) among abattoir worker in Port Harcourt, Rivers State. The difference in prevalence could be attributed to the period when the studies were embarked upon. Odinaka *et al* (2005) conducted their research in the rainy season while this study was conducted at the onset of dry season (Late November-December). The transmission of hookworm is reportedly high in rainy season (Nwosu,1981) as the eggs are commonly distributed by the rain increasing the chance of parasite-human contact Odinaka *et al* (2005). Invariably, during dry season, the transmission rate of hookworm is retarded during dry season.

The lower prevalence of *T. trichiura* recorded in this study agreed with the report of Osazuwa *et al* (2011) and Adeyeba and Tijani (2002) among children in rural communities of Edo state and malnourished school age children in peri-urban area of Ibadan respectively. Similar lower prevalence was recorded by Salawu and Ughele (2015) while Ugbomoiko and Ofoezie (2007) did not observe any *T. trichiuru* in their respective studies. However, the 13.6% prevalence observed in this study is slightly lower down the 14.9% recorded by Salawu and Ughele (2015). Although more females (54.3%) were infected than males (45.7%), there was no statistically significant (P>0.05) difference in STH infection in relation to sex. This is contrary to the record of Odinaka *et al* (2005) who reported more infection in males than in females. No clear reason could be advanced for this trend. However, this result is an indication that both sexes are vulnerable to soil transmitted helminthic infections as they both have playing contact with soil and are exposed to the same environmental, socioeconomic and hygienic conditions.

Our study showed that the only and most common co-infection was *A. lumbricoide* + *T. trichiura*. This observation is in consonance with the report of Salawu and Ughele (2015) and Oyewole *et al* (2002) who recorded a combined infection of *A. lumbricoide* + *T. trichiura* among primary school children of Ondo State, Nigeria. This result is however contrary to the records of Onuaha and Ofozie (2010) and Alli *et al* (2011). The researchers recorded co-infection of *A. lumbricoide* + Hookworm as the most common combimned infection in Nsukka, Enugu and among palm wine drinkers in Ibadan respectively.

In spite of the relatively intensity of *A. lumbricoide* in children in the age group of 5-10years (145.2epg) and 276.2epg recorded among children in Commupity primary school, Nkpor and the overall intensity of 246.5epg recorded in both school, none of the records reaches the threshold for light infection (1-4,000epg) standard spipulated by WHO (2002). Similary, the overall intensity of *T.trichura* observed in both schools (107.8epg) and that of 187epg recorded for hookworm in both schools were also below the threshold for light infection, 1,999epg and 1-1999epg for *T. trichura* and hookworm respectively WHO (2002)..

## 5. Conclusions

The results of this investigation indicate significant prevalence of STHIs in the study area, hence there is need for deliberate effort in formulate of sustainable control program by the government, Non-Governmental Organizations and other agencies including the multinational companies in the area. Provision of toilet facilities, clean drinking water, orientation on personal hygiene and improved sanitary habit will enhance control measures.

## Acknowledgments

The author wholeheartedly appreciates the assistance rendered by the head teachers, Parent Teacher Association, teachers and pupils of the schools used for this study.

REFERENCES

